# Staining of human skin with RGB trichrome unveils a proteoglycan-enriched zone in the hair dermal sheath

**DOI:** 10.1101/2022.09.20.508648

**Authors:** Clara Serrano-Garrido, Francisco Gaytán

## Abstract

The skin is the largest organ in the body and plays several essential functions acting as a barrier that protects us from physical and chemical insults, prevents the entrance of pathogens and the loss of water, besides playing an esential role in the regulation of body temperature. The skin displays a high regenerative capacity, evidenced by its self-renewing activity and the process of wound healing, driven by the existence of several resident stem cell populations. Due to the high prevalence of skin pathology, and their anatomical accessibility, skin biopsies and their subsequent microscopic observation constitute a powerfull tool for the study of primary skin diseases, as well as cutaneous manifestations of systemic diseases. This gave rise to dermatopathology as a specific discipline that unifies dermatology and pathology. In this setting, staining with hematoxylin and eosin constitutes the gold standard method for microscopic observation and diagnosis. Yet, several additional stains are used for specific purposes, such as trichrome stains for the staining of collagens in the extracellular matrix. We have applied a recently developed stain (RGB trichrome, acronym for picrosirius Red, fast Green and alcian Blue) to human samples to assess the staining outcomes in normal skin tissues. RGB staining provides a high contrasted interface between epidermis and dermis, and a comprehensible staining of the different dermal structures such as blood vessels, nerves, and sweat and sebaceous glands. The specific staining of collagens by picrosirius red can be useful for the objective quantification of these proteins under polarized light microscopy. In hair follicles, RGB staining resulted in specific staining of the epidermal sheaths and the encasing connective tissue (i.e., dermal sheath). Interestingly, the dermal sheath show two domains in which blue predominates over red staining, thus indicating that proteoglycans prevail in these areas. These two zones are the dermal papilla and an uncharacterized zone located at the upper isthmus, that we have denominated as proteoglycan-enriched belt (PEB). While the role of the papilla in the induction and regulation of hair growth is clearly established, the possible role of PEB remains to be determined and merits additional investigation.

## Introduction

The skin constitutes a large multifaceted organ that covers the external surface of the body and plays multiple functions, acting as an essential barrier to prevent pathogen entry and water loss, protects us from chemical and physical insults and plays also relevant roles in temperature regulation, sensitive interaction with the enviroment, synthesis of vitamin D, immunological defense mechanisms ^**1–3**^, and an emerging unsuspected role in the adipose tissue metabolism (mediated by sebaceous glands)^**4**^. This array of functions can be achieved due to the composition of the skin, consisting of two functionally different interacting layers (i.e., epidermis and dermis) containing several cell types and epithelial derivatives (skin appendages), that altogether conform the integumentary system.

Due to its exposed anatomical location, the skin is prone to suffer different types of injuries such as wounds or burns. Accordingly, the epidermis is constantly self-renewing and displays an outstanding regenerative capacity, that is particularly evident during wound healing^**5**^. This makes the skin a paradigm for regenerative medicine, for the study of the interactions between epithelial and mesenchymal cells, and between cells and extracellular matrix^**2**,**6**,**7**^. Among the skin appendages, hair follicles have received much attention in biomedical sciences, due to their functional and structural complexity, their self-renewing activity, their content of several stem cell populations that can regenerate the different skin cell lineages^**8–10**^, and their social relevance due to their impact in one’
ss appearance.

In general, skin diseases represent the fourth leading cause of nonfatal disease burden worldwide^**11**^, and non-melanoma skin cancer is the most common cancer affecting white-skinned people^**12**^. Due to its accessibility, microscopic examination of skin biopsies provides crucial information and constitutes the gold standard method for the diagnosis of different skin conditions. In addition to specific skin primary alterations, several systemic diseases can be detected through the study of skin histopathology. In this context, dermatopathology emerges as the combination of pathology and dermatology, and is often key for the diagnosis of skin diseases. In this setting, hematoxylin and eosin constitutes the basic routine stain for skin microscopic analysis, providing a clear nuclear and cytoplasmic differential staining. Yet, several complementary stains are available to analyze specific tissues such as connective tissue and muscle (i.e., trichrome stains) or extracellular matrix components such as proteoglycans (i.e., alcian blue). A recently developped trichrome stain (RGB trichrome)^**13**^ is composed of two primary dyes (picrosirius red and alcian blue, in addition to fast green) that specifically stain two of the main components of the extracellular matrix, i.e., collagen and proteoglycans. Given the complex structure of the skin and adnexial appendages, consisting of a mixture of both epithelial and mesenchymal tissues, it is worth assessing if RGB staining provides valuable information when applied as a complementary stain to the routine H&E for microscopic examination. This could be useful for skin research and pathology, and is potentially applicable to tissue segmentation and automated image analysis^**14**^.

## Materials and Methods

Human skin samples were obtained from the Bio-archive of the Department of Pathology of the Faculty of Medicine and Nursing. These specimens corresponded to skin biopsies that were obtained for diagnostic purposes due to the presence of different skin lessions. Areas of the skin at the margins of the biopsy, that were free of pathology, were used for this study. These skin samples were collected between 1980 and 2000, in keeping with contemporary legislation, including informed consent. Anyway, the use of these samples was approved by responsible members of the Department of Pathology, granting patient confidentiality and following strict adhesion to current legislation.

Human tissues had been fixed in 4% phosphate buffered formaldehyde and embedded in paraffin. A total number of 50 skin samples were studied. Six μm-thick sections were cut from paraffin blocks, and after dewaxing and rehydration were submitted to RGB staining, following previously described methods^**13**^. Briefly, the sections were sequentially stained with 1% (w/v) alcian blue in 3% aqueous acetic acid solution (pH 2.5) for 20 min, washed in tap water, stained with 0.04% (w/v) fast green in distilled water for 20 min, washed in tap water, and finally stained with 0.1% (w/v) sirius red in saturated aqueous solution of picric acid for 30 min, washed in two changes of acidified (1% acetic acid) tap water, dehydrated in 100% ethanol, cleared in xylene, and mounted with synthetic resin. Some human skin sections from the files of the Department of Pathology, that were already stained with hematoxylin and eosin or Masson trichrome, were used (as indicated in figure legends), in order to compare them with RGB-stained tissues.

## Results

The structural features and functional properties of the integumentary system (skin and skin appendages) have been clearly established and described^**1**,**2**,**15**^. Here, we will made a brief description, limited to those tissue structures exhibiting relevant outcomes after staining with RGB trichrome. Special emphasis will be devoted to the staining characteristics of the hair follicle, as it constitutes an interdisciplinary model for biomedical research^**9**,**10**,**16**^. All figures in this study are stained with RGB trichrome unless otherwise indicated in the figure legend. Scale bars have been included as a reference in some panels per figure.

### General staining of the skin

Human skin displays significant regional differences troughout the body^**17**^. However, the staining properties of the skin are not dependent on its topographical origin, except for the relevant differences that exist between thin (covering most of the body surface) and thick (located mainly in palms and soles) skin types. In all cases, staining of the human skin with RGB trichrome allows a clear differentiation between epidermis and dermis. Whereas the epidermis is pale-stained with green-stained keratin in the stratum corneum, the dermis contains abundant collagen and is intensely red stained (Fig. 1). Keratinocytes in the epidermis appear as pale grenish-stained polyhedral cells that are clearly appreciated in the stratum spinosum (Fig.1C). In the dermis, several structures such as hair follicles, blood vessels and sensitive copuscles are present (Fig 1E-G). Differences in the thickness of the collagen bundles exist between the papillary and reticular dermis (Fig. 1B,D). The ducts of dermal glands make their way to the skin surface through the epidermis (Fig. 1H). In glabrous (thick) skin, five different epidermal layers can be appreciated (Fig. 2). The stratum granulosum stains red, whereas the stratum lucidum is green stained and the stratum corneum show a mixture of green and red stained areas (Fig. 2A-C). In the abundant dermal papillae, blood capillary loops and Meissner corpuscles can be observed (Figs. 2 D,E). The corkscrew-shaped ducts of the sweat glands pass through the stratum corneum (Fig. 2F).

**Figure 1.**
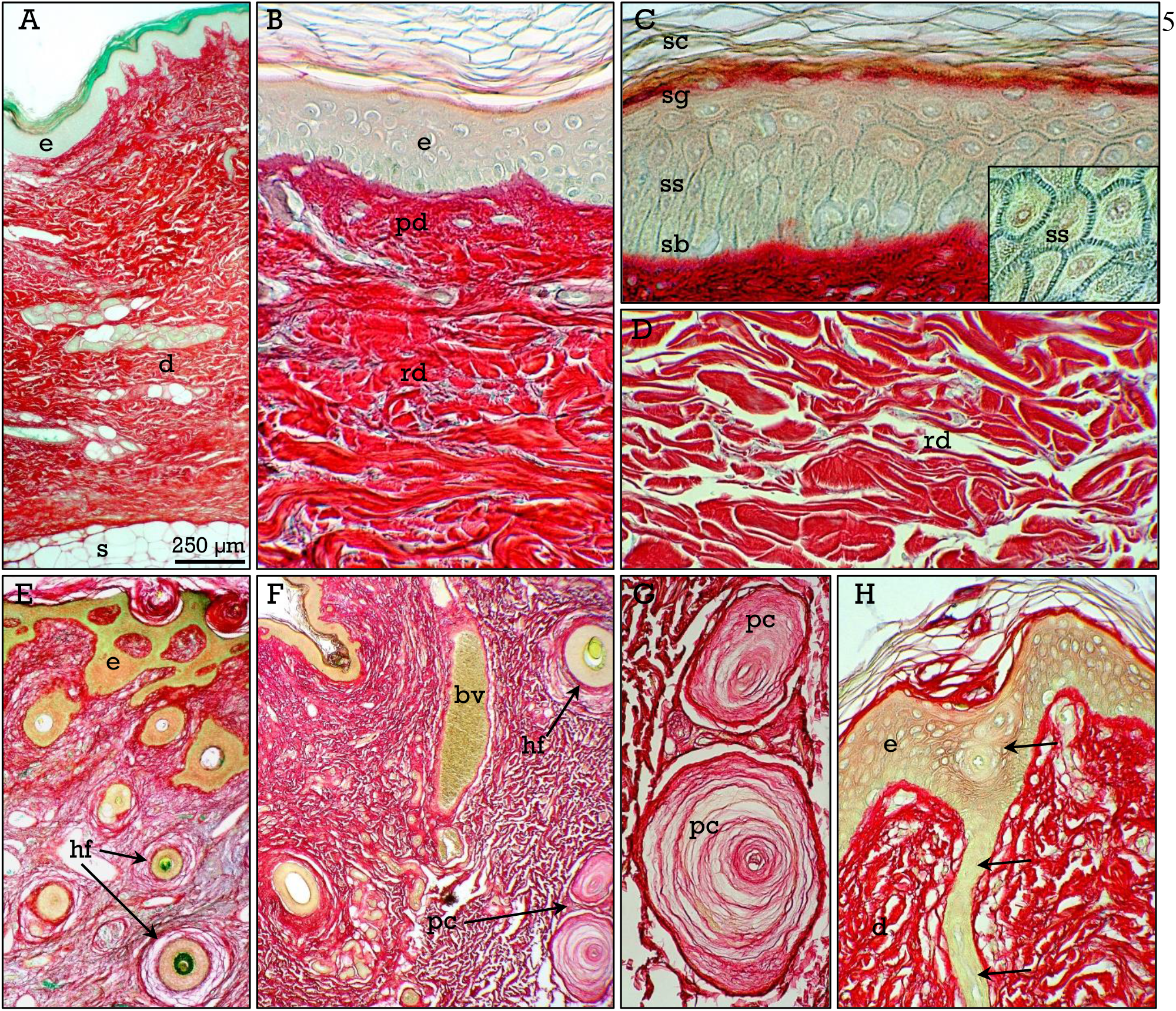
Thin skin. The skin (**A**) is composed of an epithelial and a mesenchymal layer, the epidermis (***e***) and the dermis (***d***), resting on the hypodermis or subcutaneous tissue (***s***). The epidermis (**A-C**) consist of a stratified keratinized epithelium in which several layers can be distinguished: stratum basalis (***sb***), spinosum (***ss*** and ***inset***), granulosum (***sg***) and corneum (***sc***). The dermis (**A**,**B**,**D**) can be divided into two zones, the papillary (***pd***) and the reticular (***rd***) dermis, containing collagen bundles, that are responsible for the intense red staining, and are thicker in the reticular layer (**D**). In the dermis, different structures such as hair follicles (***hf***), blood vessels (***bv***), and sensitive pacinian corpuscles (***pc***, shown at higher magnification in **G**), can be observed (**E-G**). The ducts of the dermal glands (***arrows***) make their way to the skin surface through the epidermis (**H**).

**Figure 2.**
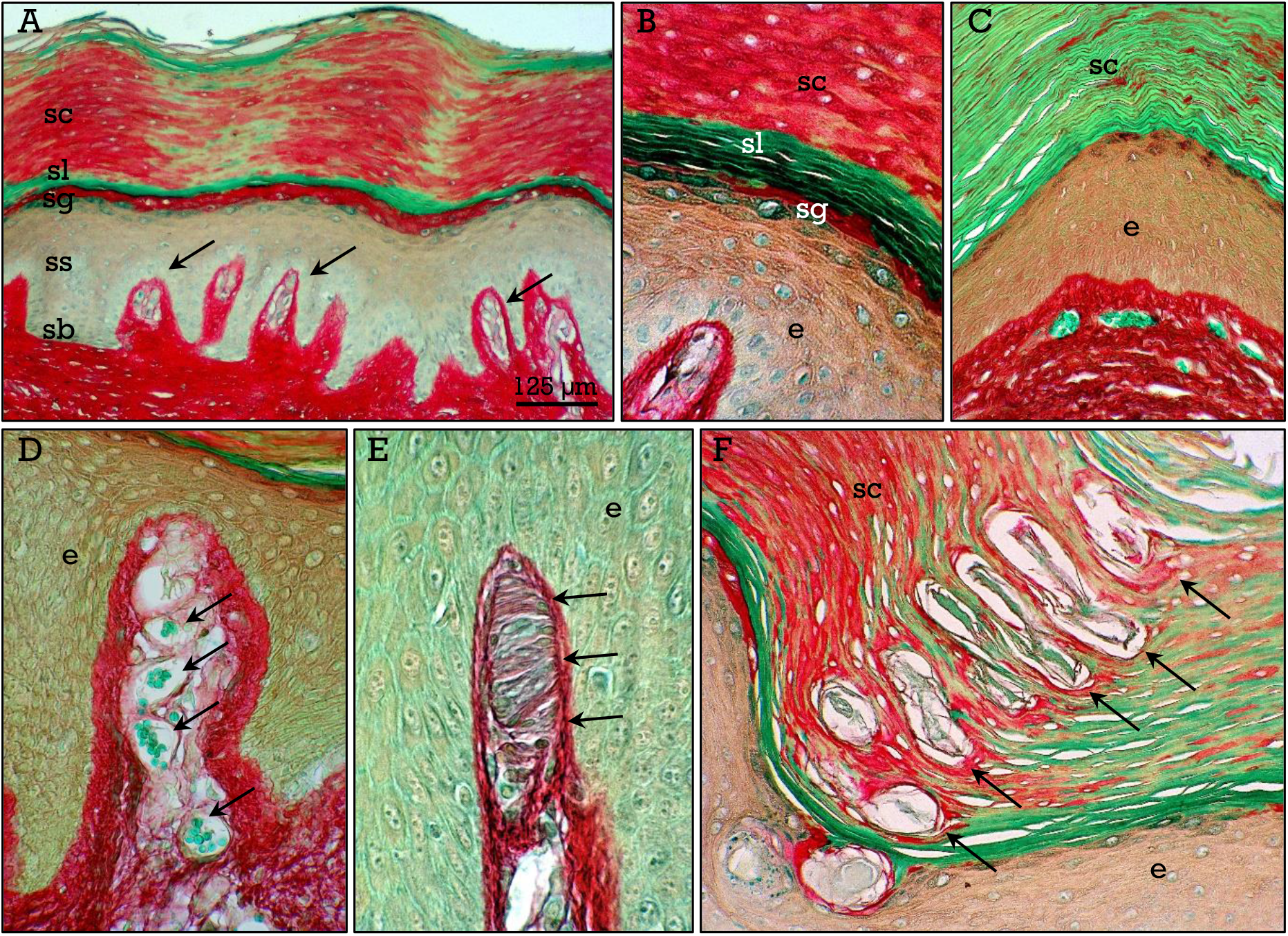
Thick skin. The skin (**A**) in the palms and soles has a thick keratinized epidermis in which five different layers, stratum basalis (***sb***), spinosum (***ss***), granulosum (***sg***), lucidum (***sl***), and corneum (***sc***) can be distinguished. The stratum lucidum stain green whereas the stratum corneum (***sc***) shows a mixture of red and green stained areas (**A-C**). The dermis show abundant papillae (***arrows*** in **A**) in which capillary loops (***arrows*** in **D**) and Meissner corpuscles (***arrows*** in **E**) can be observed. The sudoriparous glands open on the surface by a corkscrewlike duct through the stratum corneum (***arrows*** in **F**).

In addition to keratinocytes, other cell types such as melanocytes and Langerhans cells, among others, are present in the epidermis (Fig. 3). Melanocytes appears as small clear cells intermixed with the basal keratinocytes (Fig. 3A). However, when melanocytes are located in the dermis (intradermal nevus; Fig. 3B) or in the hair follicle bulb (Fig. 3C), melanin accumulates in the cytoplasm and appear as brown/black cells. Melanin can be also appreciated in a supranuclear location in basal epidermal keratinocytes (Fig. 3D) in people with brown of tanned skin. Otherwise, Langerhans cells appear as clear cells scattered between the keratinocytes in the stratum spinosum (Fig. 3E). Both melanocytes and Langerhans cells can be detected by immunostaining for S100 protein (Fig. 3F)^**18**^. Interestingly, counterstaining with RGB trichrome does not interfere with the diaminobenzydine-derived chromogen. In the dermis, blue-stained mast cells (Fig. 3G) can be observed. Otherwise, mast cells (Fig. 3G) can be observed in the dermis, whereas adipocytes are the main component of the hypodermal (i.e, subcutaneous) tissue (Fig. 3H).

**Figure 3.**
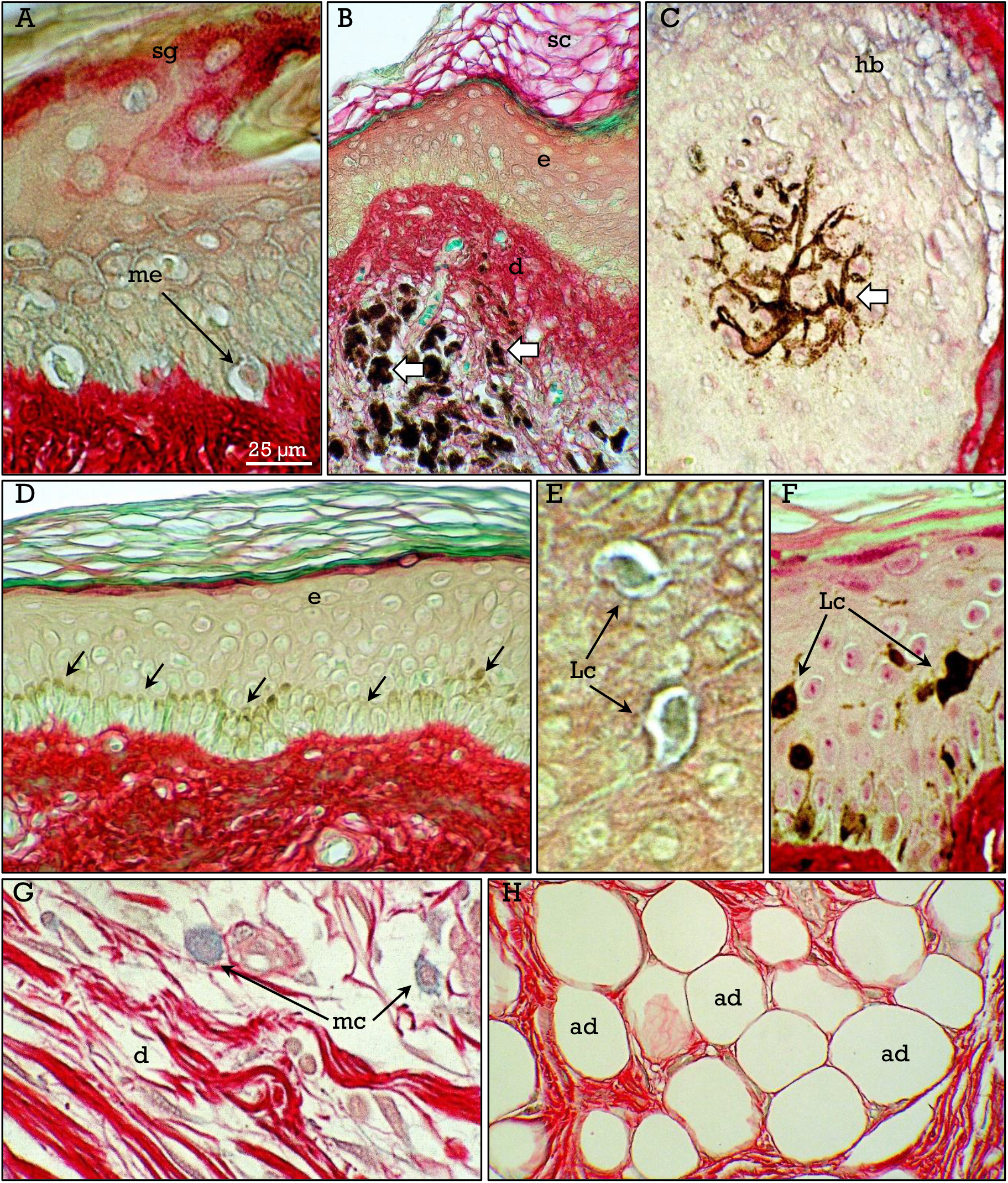
Other skin cell types. In the epidermis, melanocytes (**A-C**) appears as clear cells at the base of the epithelium (***me*** in **A**), and as brown-stained cells in a blue nevus located in the dermis or in the hair follicle bulb (***arrows*** in **B** and **C**). Melanin can also be seen in the supranuclear zone of the basal keratinocytes (***arrows*** in **D**) in pigmented skin. Langerhans cells appears as clear cells scattered between keratinocytes (***Lc*** in **E**). In **F**, skin section immunostained for S100 protein that labels Langerhans cells, counterstained with RGB trichrome. Mast cells are present in the dermis (**G**), and adipocytes (***ad***) in the subcutaneous tissue (**H**).

### Sweat and sebaceous glands

Practically all histological tissues are present in the skin. The dermis contains different structures such as blood vessels, nerves, and three types of glands (Figs. 4,5) that are responsible for the nutrition of the skin and for several specific functions that make the skin the largest sensitive organ, and the main regulator of body temperature. Myelinated nerves can be easily identified in both transverse and logitudinal sections (Fig. 4A-C). The three types of glands correspond to eccrine and apocrine sweat glands and sebaceous glands. Eccrine sweat glands are distributed throughout the skin^**3**,**19**^, and are particularly abundant in the thick skin of the palms and soles. These glands appear as pale green-stained tubules (Fig. 4D-F) reaching the deep dermis. The secretory and duct zones can be distinguised by their different shades of green color (Fig. 4D,E). Although eccrine sweat glands open independently on the skin surface, recent studies have reported a close relation between them and the pilosebaceous units (Fig. 4F). Apocrine sweat glands are restricted to specific areas such as the axillae and the perineal zone, and are associated to pilosebaceous units^**3**^. These glands show intensely green stained secretory epithelium, composed of a layer of cells showing apical protrusions and a dilated lumen (Fig. 4G,H). The apical zones of epithelial cells and the milky secretory product, that accumulates in the duct of some glands, are intensely blue-stained (Fig. 4H,I). Finally, sebaceous glands are associated to hair follicles and located in the mid dermis, and are composed of pale blue-stained vacuolated cells (Fig.5). These glands are present throughout the body except in some specific areas such as the glabrous skin of palms and soles.

**Figure 4.**
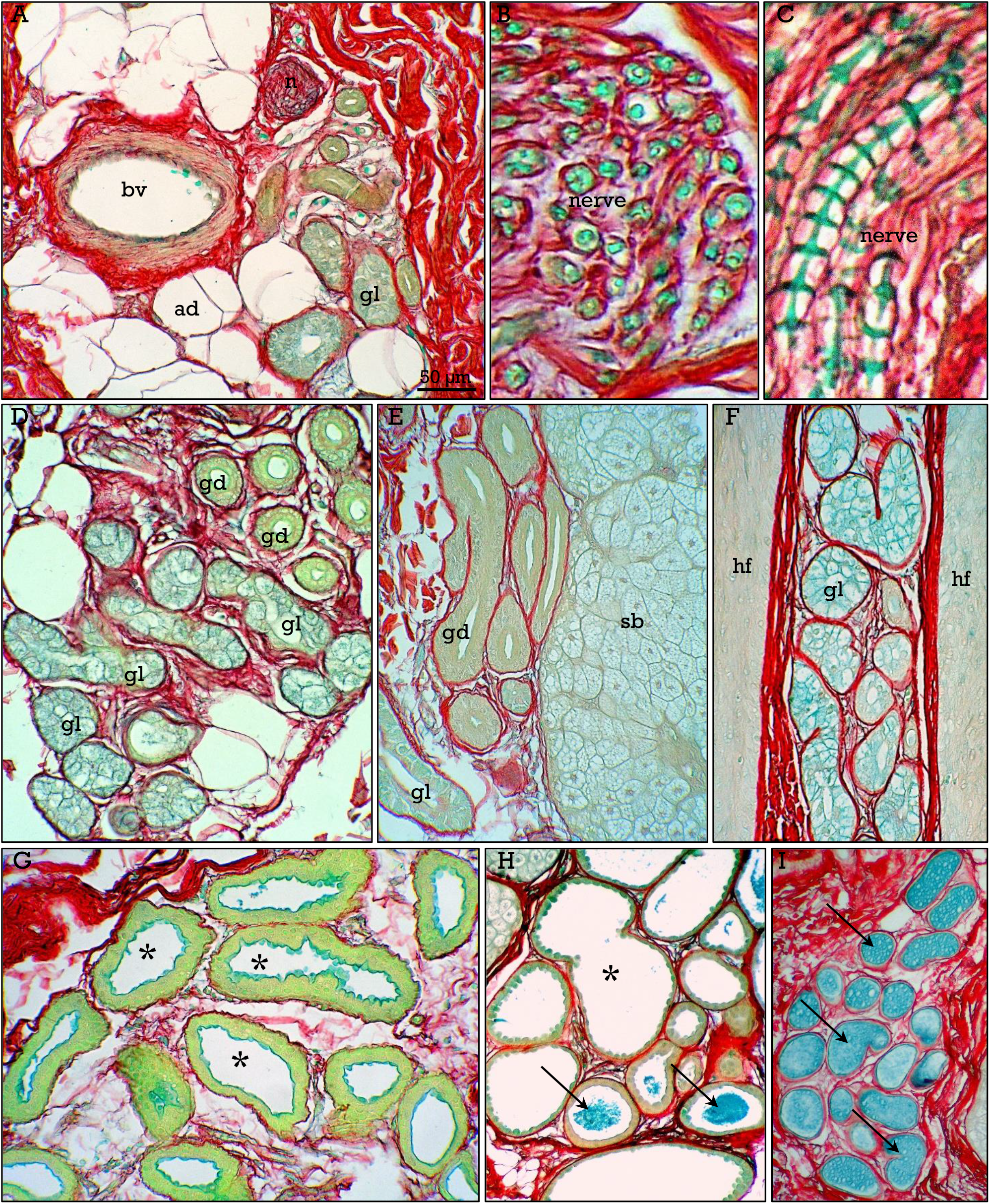
Dermal structures. Sweat glands. The dermis (**A**) contains blood vessels (***bv***), nerves (***n***) glands (***gl***) and some adipocytes (***ad***). Nerves are clearly distinguised in either transverse (**B**) or longitudinal (**C**) sections. In the later, the Schmidt-Lanterman incisures can be observed. Eccrine sweat glands (**D-F**) are pale green-stained. Secretory portion (***gl***) and ducts (***gd***) showed slightly different shades of color (**D**,**E**). In some cases, eccrine sweat glands are related to sebaceous glands (***sb*** in **E**) or hair follicles (***hf*** in **F**). Apocrine sweat glands (**G-I**) showed dilated lumen (***asterisks***) with an intensely green-stained epithelium, contrasting with the blue-stained secretory material that can be observed at the apical zone of epithelial cells and accumulated in the gland ducts (***arrows*** in **H, I**).

**Figure 5.**
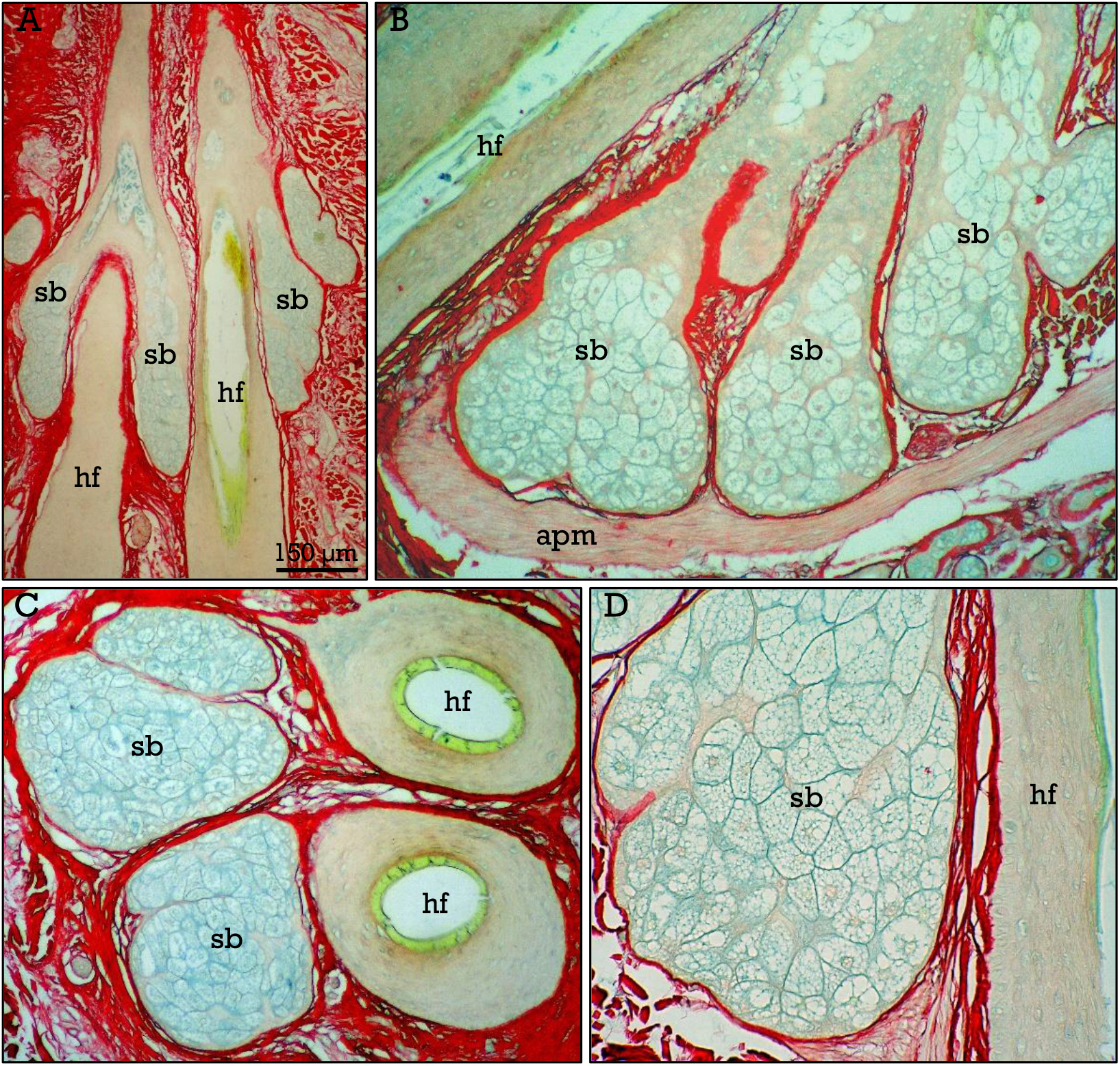
Sebaceous glands. Sebaceous glands (***sb***) are associated (**A-D**) to hair follicles (***hf***), located in the mid dermis, showing a vacuolated pale blue-stained cytoplasm, and are closely related (**C**) to the arrector pili muscle (***apm***).

### Hair follicles

Hair follicles, which form the pilosebaceous units together with the sebaceous glands and the arrector pili muscle (APM), are skin appendage miniorgans that show a high structural and functional complexity^**15**,**16**^. Hair follicles self-renew periodically, alternating growth and resting phases, during which the lower segment undergoes important structural changes, regressing during catagen, resting during telogen, and undergoing new growth activity during anagen phase (Figs. 6,7), whereas the upper segment remains practically unchanged^**16**,**20–22**^. There are two types of hair follicles. Vellus hairs are fine and non-pigmented and covers most of the body except some specific areas such as the palms and soles, and their bulbs are located in the dermis. Terminal hairs are present in the scalp, axillae, and in beard, chest, legs and arms in males. They are thicker, pigmented, and anchored in the subcutaneous tissue^**15**^.

**Figure 6.**
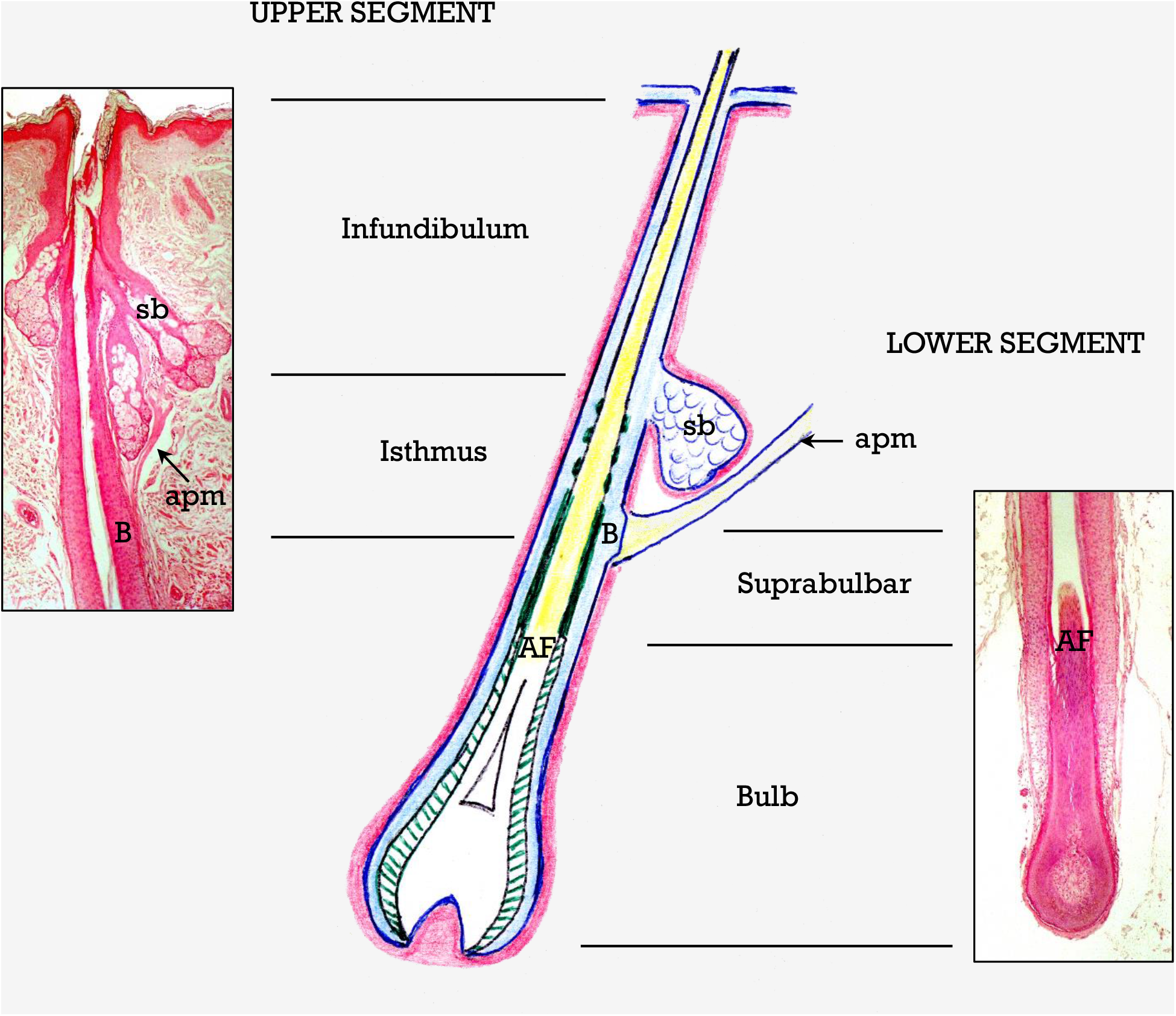
Hair follicle segments. Schematic drawing depicting the different zones of the upper and lower segments of the hair follicle. Micrographs correspond to H&E stained follicles. ***sb***, sebaceous glands; ***apm***, arrector pili muscle; ***B***, bulge area; ***AF***, Adamson fringe.

**Figure 7.**
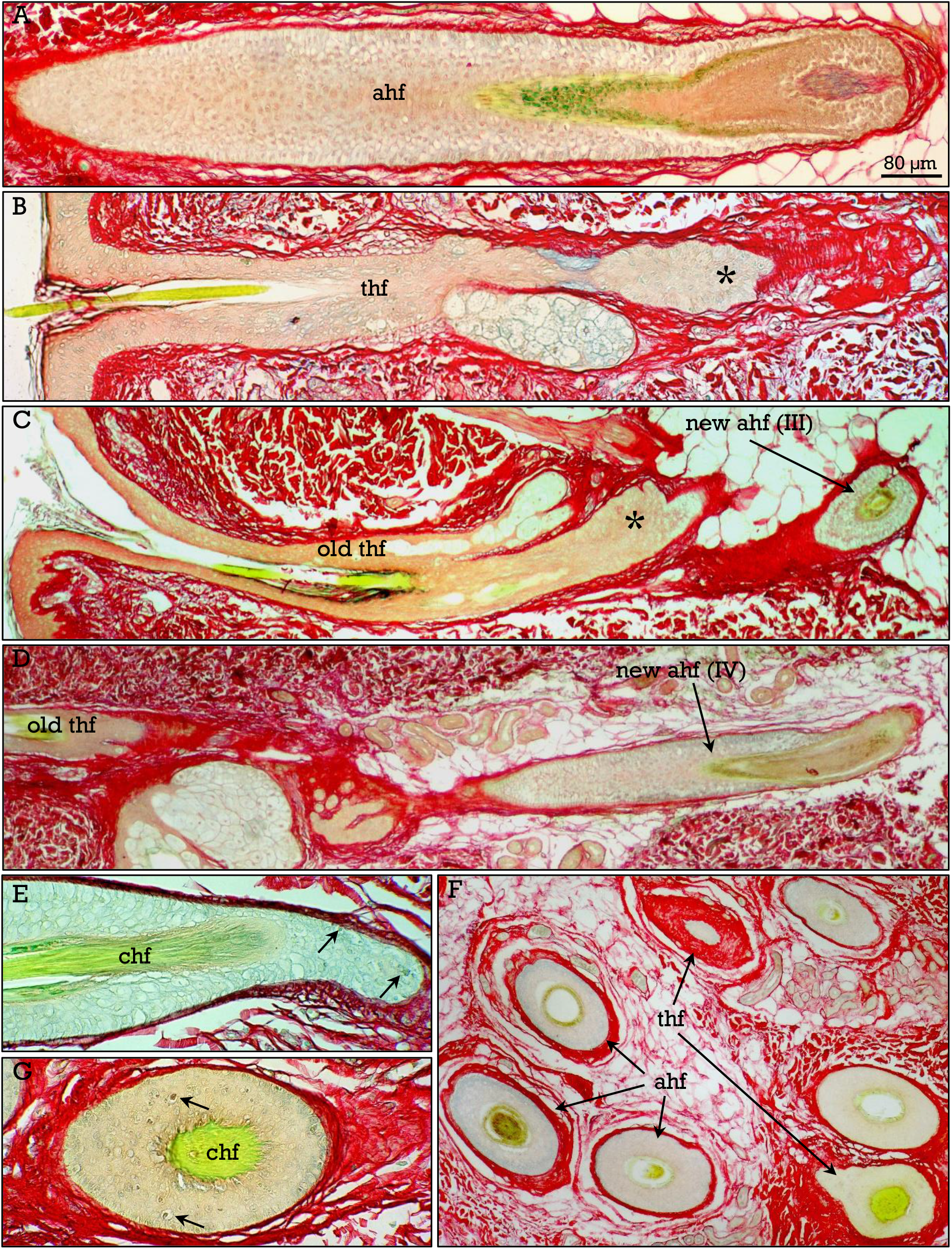
Hair follicle cycle. Terminal follicles in anagen (**ahf**), telogen (**thf**) and new growth anagen III (**C**) and IV (**D**) phases. **E, F** hair follicles in catagen (***chf***), with several apoptotic cells (***arrows***). **G**, transverse sections of hair follicles in anagen and telogen.

A schematic drawing depicting hair follicle anatomy is shown in Figure 6. The hair follicle can be divided into two (upper and lower) segments. The upper segment consists of the *infundibulum* (from the ostium to the entrance of the sebaceous duct) and the *isthmus*, from the duct of the sebaceous glands to the site of attachment of the APM. The lower segment consists of the *suprabulbar zone*, from the insertion of the APM to the Adamson’
ss fringe (AF), and the *bulb* (below the AF) containing the papilla and the matrix cells. The hair follicles are composed of several epithelial and mesodermal components that undergo morphological changes from the bulb to the ostium at the skin surface. Details of the hair follicle cycle are shown in Fig. 7, and the structure of anagen hair follicle is shown in Fig. 8. The hair shaft is composed of the cuticle, and the cortex that is fully keratinized and anucleated above the AF. The hair shaft is surrounded by two concentric epithelial sheaths. The inner root sheath (IRS) consists of three layers that are, from inside to outside, the cuticle, the Huxley, and the Henle layers. At the upper isthmus the IRS that is fully keratinized breaks up and is lost. The outer root sheath (ORS) is an invagination of the epidermal epithelium and extend from the bulb to the skin surface, with no evident morphological changes. At the insertion of the APM there is a small protuberance that constitutes the bulge. In the infundibulum the ORS is the only sheath surrounding the hair shaft.

**Figure 8.**
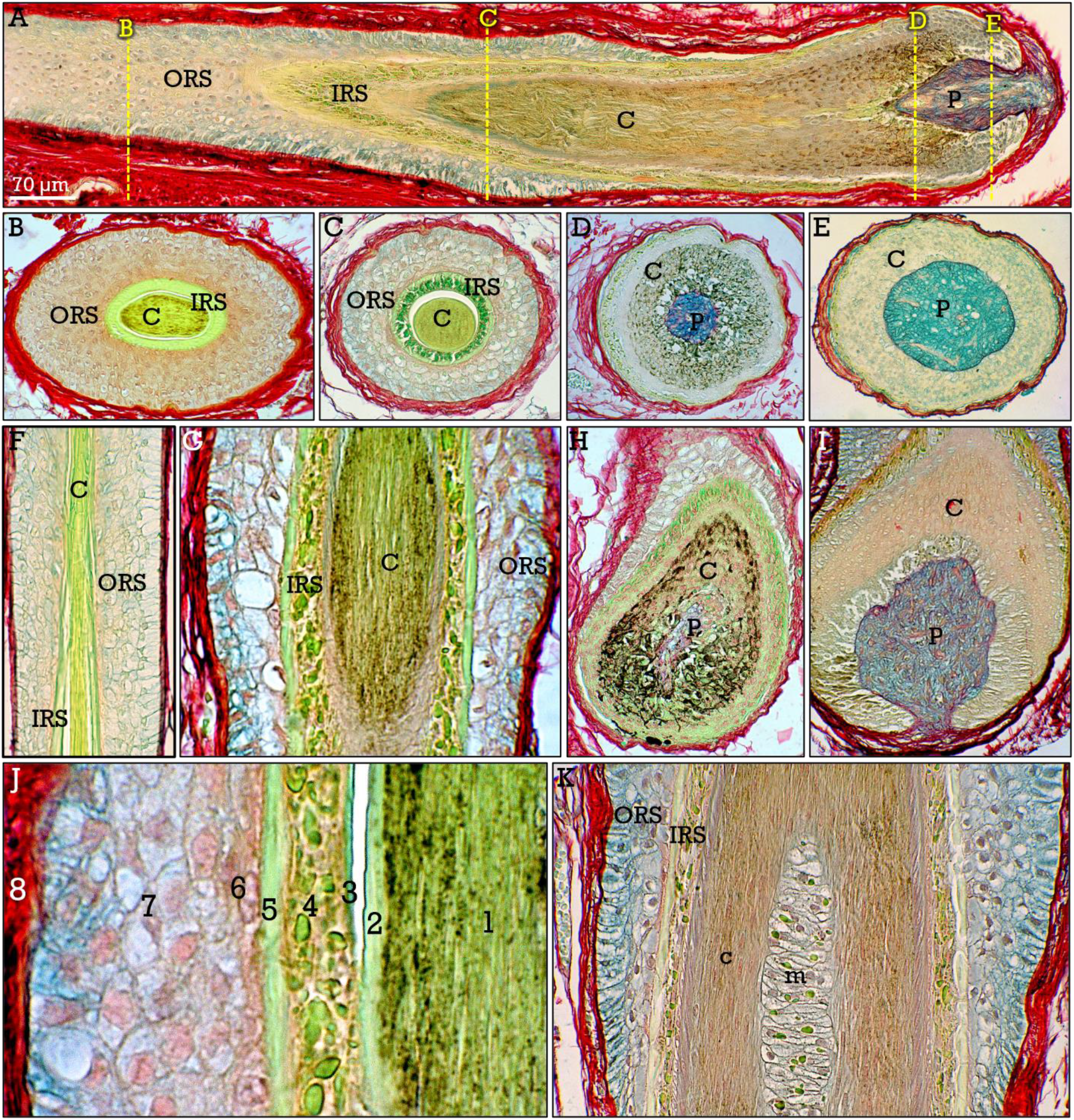
Structure of the hair follicle. Transverse (**B-E**) and longitudinal (**F-I**) sections at different levels (indicated in **A**) of the hair follicle. The different structures, papilla (***p***), cortex (***c***), inner root (***IRS***) and outer root (***ORS***) sheaths are indicated. **J**,**K**, longitudinal sections, showing the different layers: hair medulla (***m***), hair cortex (***1***), hair cuticle (***2***), IRS cuticle (***3***), Huxley (***4***) and Henle (***5***) layers, companion layer (***6***), ORS (***7***), and dermal sheath (***8***).

The staining properties and morphological features of the hair follicle at different levels from the bulb to the infundibulum are shown in transverse and longitudinal sections in Figs. 8–10. Sections at the lower bulb level show the blue-stained dermal papilla (Fig. 8A,E,I). At a higher level in the bulb the melanization of the matrix cells is evident (Fig. 8A,D,H). Sections at the upper bulb, show the green-stained IRS with abundant trichohyaline granules, and the ORS (Figs. 8A,C,G), and at the suprabulbar zone, the ORS and the keratinized IRS (Fig. 8A,B,F) can be observed. The different layers of the IRS^**23**^ and ORS can be identified (Fig. 8J). In some terminal hairs, the central medulla is present (Fig. 8K).

The Adamson fringe (AF)^**24**^ constitutes a zone at which important changes in the hair shaft and IRS (Fig. 9) occur. So, at this level the hair cortex cells loss their nuclei and become fully keratinized, and the Huxley layer of the IRS loss trichokeratin granules and keratinizes, while the Henle layer is already keratinized at the upper bulb. Consequently, the whole IRS is keratinized above the AF at the suprabulbar region. Staining with RGB trichrome evidenced that some histochemical changes also occur in the ORS at the AF level, although morphological differences are not evident. The ORS located at the bulb (i.e., below the AF) stains pale blue (Figs. 9A, 10C), but this staining vanishes at the suprabulbar region (i.e., above the AF; Figs. 9A, 10E). This shift in ORS staining properties is fully coincident with the location of the AF, as evidenced in oblique sections at this level (Fig. 10A,D). These staining properties of the ORS were mainly observed in terminal hair follicles whereas blue staining was weak or barely detectable in some vellus hairs. Another staining characteristic of the ORS that change along the follicle is PAS+ staining, due to the accumulation of glycogen^**25**^. Although PAS positivity increases in general toward the bulb, it is not fully coincident with changes in RGB staining, since PAS+ staining in the ORS is not restricted to the bulbar region (i.e., delimited by the AF), and can be observed both below and above the AF level (Fig. 9A). The inner cells of the ORS, in contact with the Henle layer of the IRS (Fig. 8J), constitute the companion layer^**26**^.

**Figure 9.**
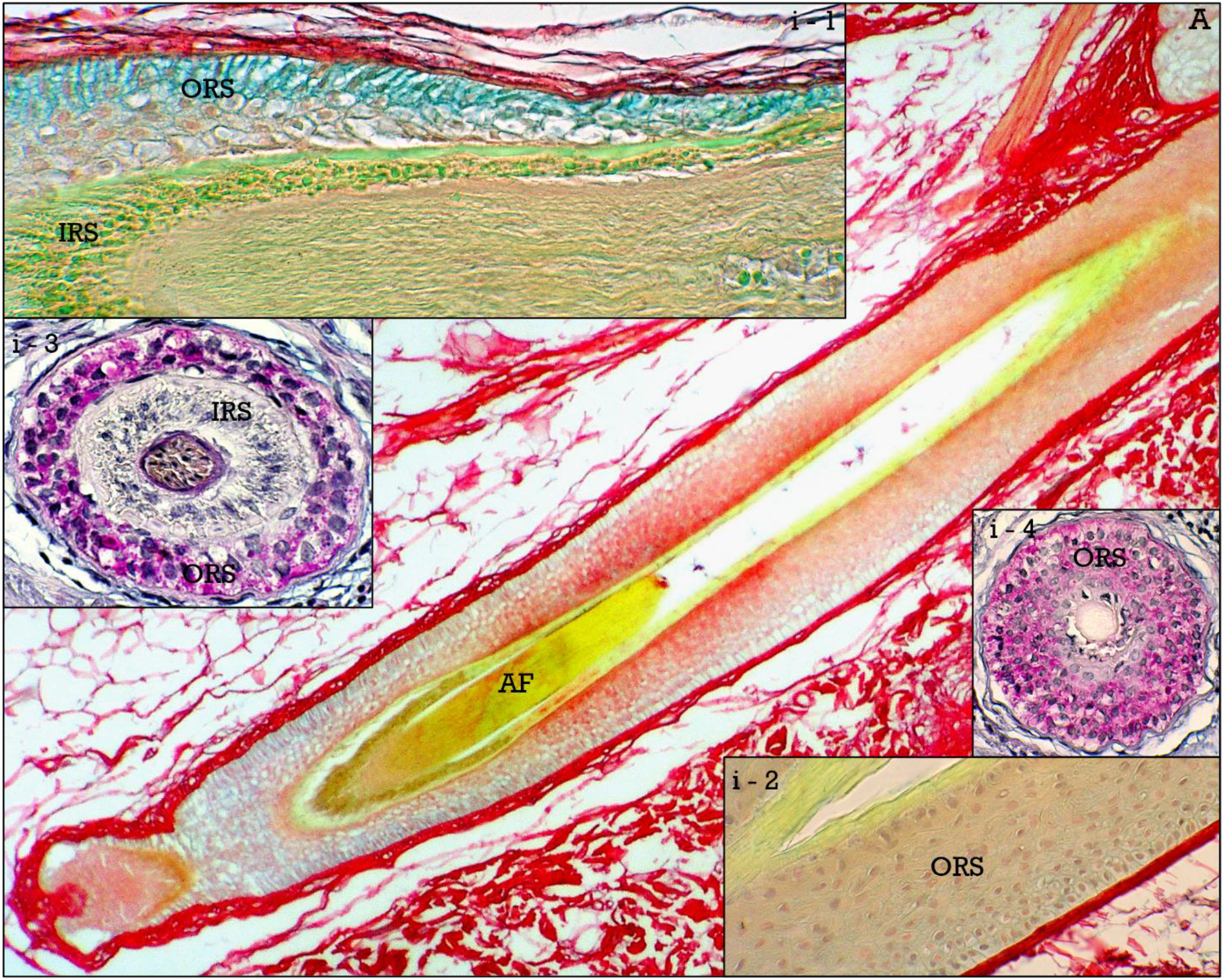
Outer root sheath (ORS). **A**, shift in ORS staining at the Adamson fringe (***AF***) level. At the bulb, below the AF, the ORS stains pale blue (***i1***) but this color vanishes in the suprabulbar zone (***i2***). The ORS is PAS+ either above (***i3***) or below (***i4***) the AF.

**Figure 10.**
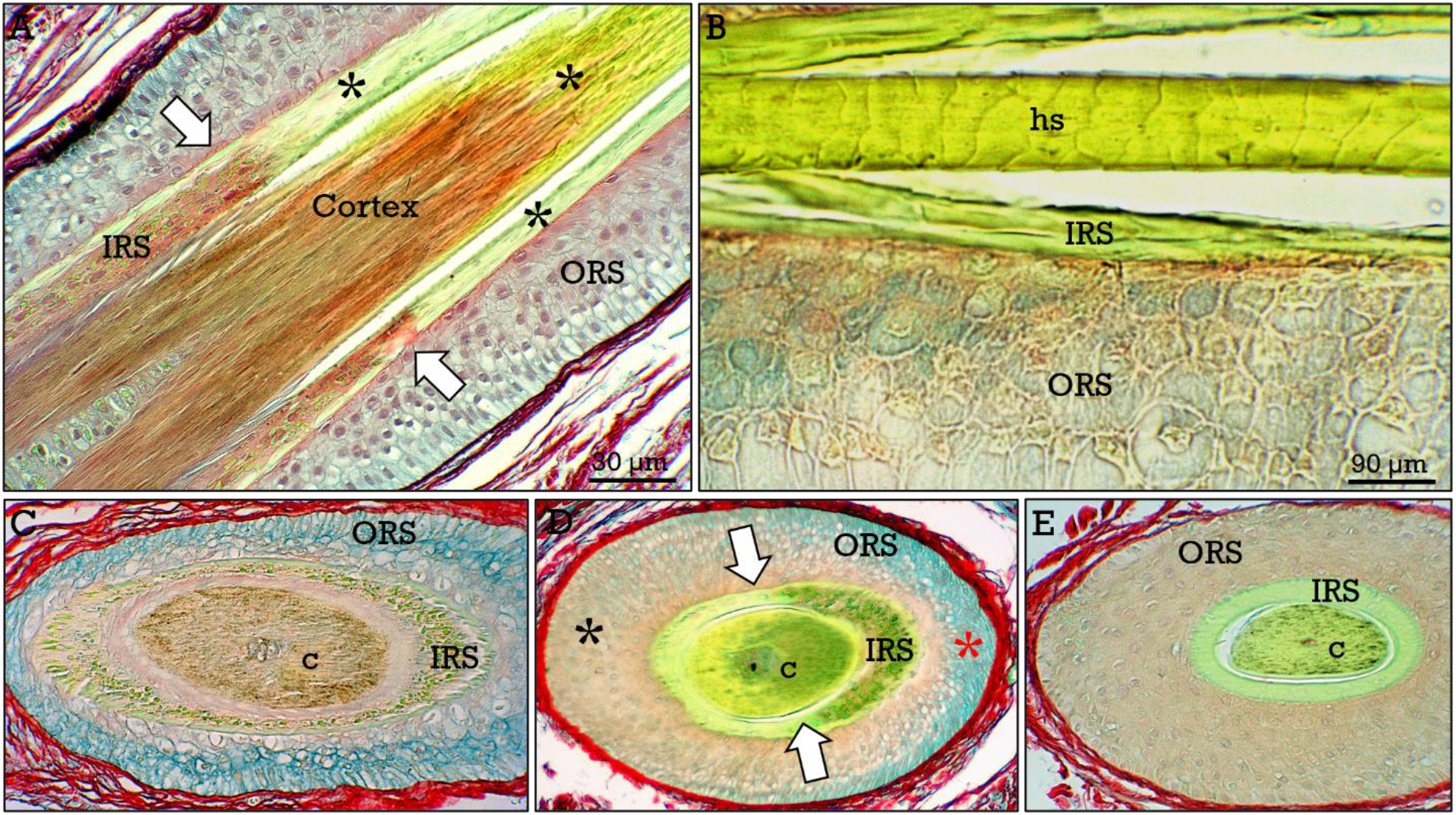
Adamson fringe. The AF area (***arrows***) is shown in **A**. At this level, the hair cortex and the IRS become keratinized (***asterisks***), as can be observed in longitudinal (**C**) sections above the AF. The ORS is shown in transverse sections below (**D**), and above (**F**) the AF. In slightly oblique sections (**D**) at the AF level, the shift in the staining of the ORS can be observed (***red*** and ***black asterisks***, just above and below the AF respectively).

In addition to the concentrical epithelial layers of the hair follicle, the dermal sheath is a layer of connective tissue encasing the whole follicle^**27**^, which is stained red due to its high collagen content (Fig.11). This is separated from the ORS by a basement membrane (aka vitreus membrane; Fig. 11B). The dermal sheath have specialized domains such as the dermal papilla^**28**,**29**^ and the zone of insertion of the APM (bulge area)^**30**^. The papilla is formed by connective tissue composed of blood vessels and a cluster of mesenchymal cells, and stains blue with RGB trichrome (Fig. 11A). In some areas, myelinized nerves can be observed in the dermal sheath around the hair follicle (Fig. 11F). The bulge area (Fig. 12) located at the insertion of the arrector pili muscle, is particularly prominent in telogen hair follicles (Fig. 12A-C), and seems to be more subtle in anagen follicles (Fig. 12E,F). This region is highly relevant by the presence of a population of hair associated stem cells^**31–33**^.

**Figure 11.**
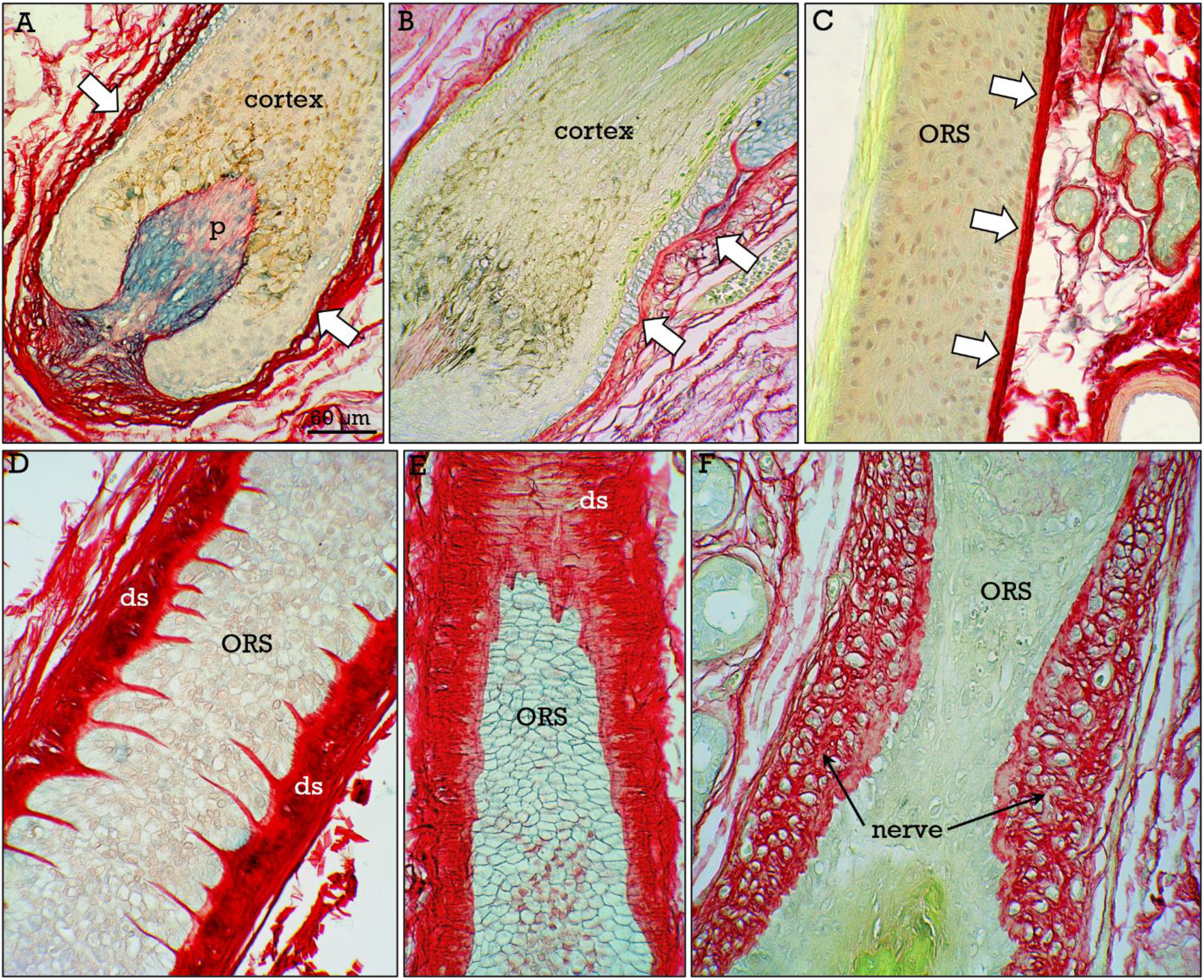
Dermal sheath. The dermal sheath (***arrows***) consists of the papilla (***p*** in **A**), and the connective tissue layer encasing the hair follicle (**B-F**). The dermal sheath is separated from the ORS by the vitreus membrane (***arrows*** in **B**). In tangential sections (**D**) the dermal sheath-ORS interface (**D**) and the external aspect of the ORS (**E**) can be appreciated. Nerves surrounding the hair follicle are shown in **F**.

**Figure 12.**
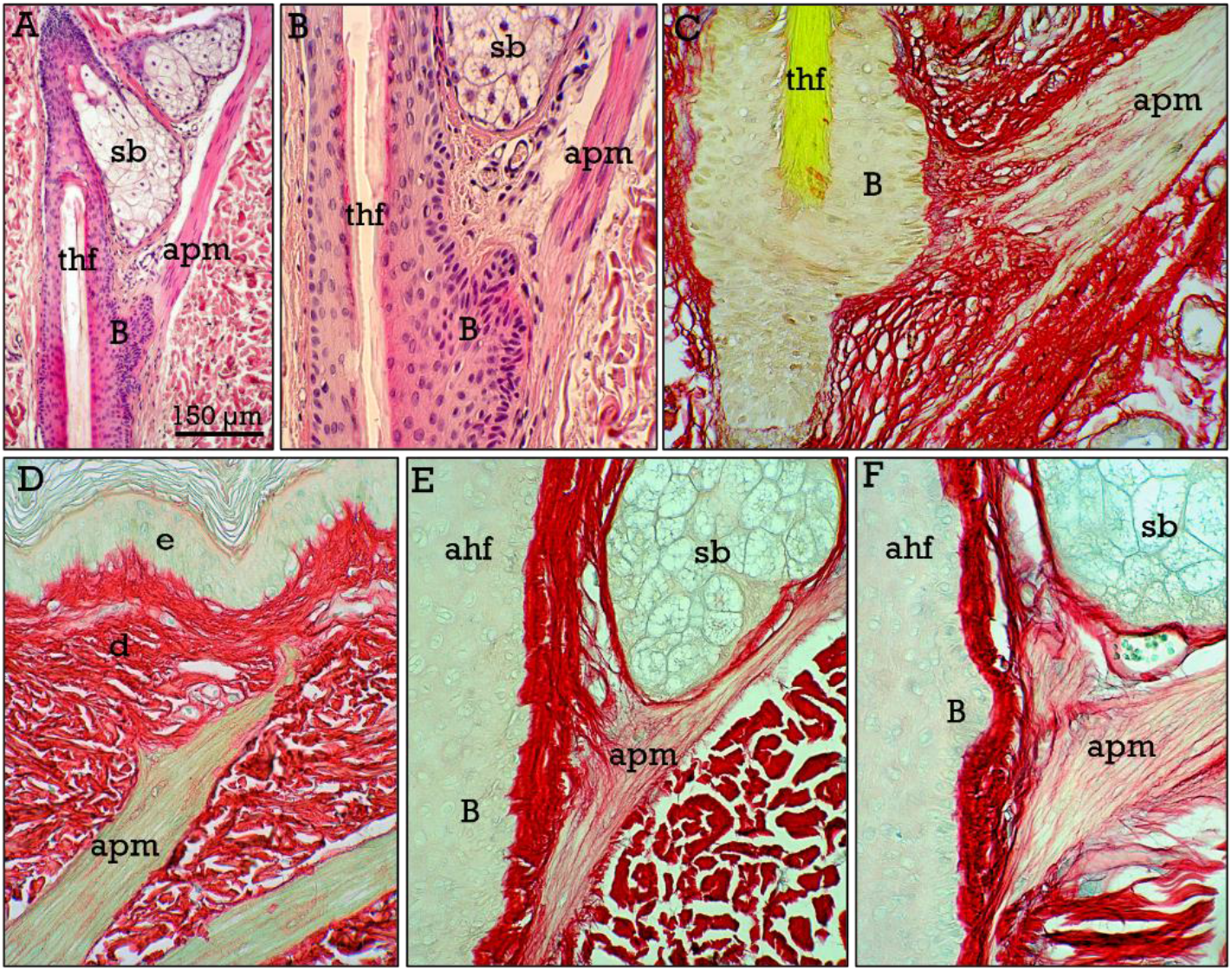
Bulge area. The bulge area is shown in H&E (**A**,**B**) and RGB trichrome (**C-I**) stained sections. The bulge (***B***) is located at the point of insertion of the arrector pilli muscle (***apm***) into the dermal sheath of the hair follicle (***hf***). The apm is also attached to the papillary dermis (**D**). The bulge seems to be more prominent in telogen (***thf*** in **A-C**) than in anagen (***ahf*** in **E**,**F**) follicles.

One of the most relevant features of the RGB stained hair follicles is the differential staining of a zone of the dermal sheath located at the upper isthmus, just below the junction to the sebaceous gland duct. This zone forms a belt of blue-stained extracellular matrix around the ORS (Figs., 13, 14), that we have denominated as PEB (acronym for Proteoglycan-Enriched Belt). In logitudinal sections (Fig. 13), the PEB appears as two lateral bands, in which the collagen (red-stained) was scarce and the dermal sheath turns blue (i.e., alcian blue-stained), strongly suggesting that proteoglycans are the predominant component of the dermal sheath in this zone. The belt-shaped PEB can be better appreciated in transverse sections of the hair follicle (Fig. 14). Furthermore, in these sections, the precise location of the PEB in the hair follicle can be determined, taking as reference the structural changes in the IRS (Fig. 14). The PEB is located in the zone where the fully keratinized IRS is fragmented or just shed, whereas is absent in the zone where the IRS is unkeratinized or at least not (or initially) fragmented (Fig. 14A,E). From this point, the PEB extends up to the junction of the sebaceous gland duct, and is not found in the infundibulum (Fig. 14F). The distinction between vellus and terminal hair follicles can be also determined in transverse sections, as in the former, the hair shaft diameter is equal or less than the thickness of the IRS^**15**^. In general, the PEB seems to be more prominent in vellus than in terminal hairs. In terminal hairs of the scalp, only small blue-stained areas can be appreciated (Fig. 13E-G). At high magnification, these areas contain collagen bundles and scattered cells in a blue-stained matrix (Fig. 13E,G), and myelinated nerves were also observed in this area in some follicles (Fig.13F). Otherwise, the possible changes of the PEB along the different phases of the hair follicle cycle were not investigated.

**Figure 13.**
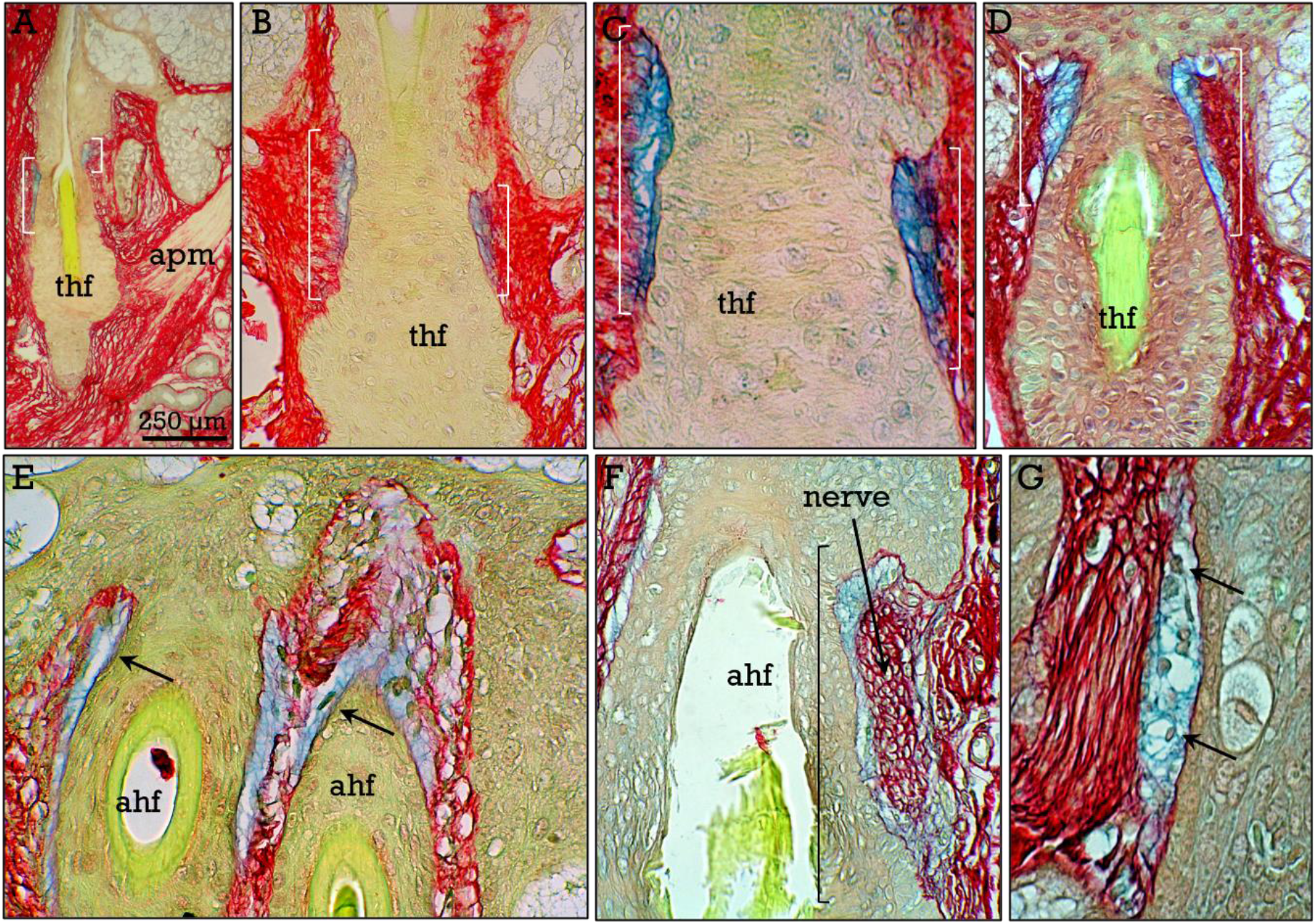
Proteoglycan-Enriched Belt (PEB). Longitudinal sections showing the PEB (delimited by ***brackets***) in telogen (***thf***), and anagen (***ahf***) hair follicles. Mielinated nerves (**F**) and scattered cells (**G**) are present in the PEB in some follicles.

**Figure 14.**
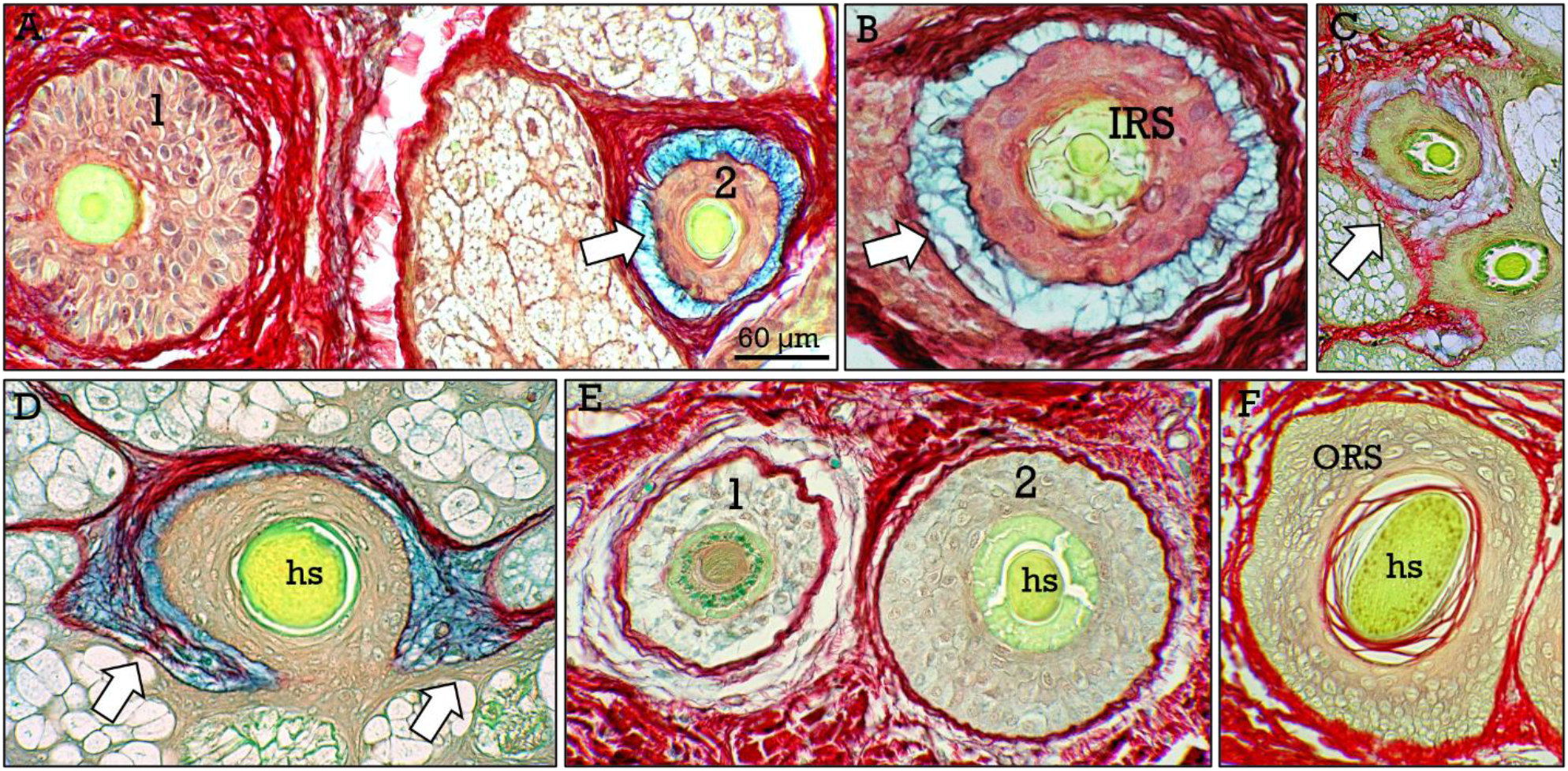
PEB in transverse hair follicle sections. The belt-shaped PEB (***arrows***) can be better appreciated in transverse sections. In **A**, the PEB is present in the follicle lacking IRS (***2***), but not in the vellus hair with an intact keratinized IRS (***1***). Similarly, the PEB can be observed in hair follicles with fragmented (**B**,**C**) or absent (**D**) IRS, but not (**E**) in those showing a non keratinized (***1***) or initially fragmented (***2***) IRS. The PEB is also absent in the infundibulum (**F**). ***sb***, sebaceous glands; ***hs***, hair shaft.

## Discussion

Skin biopsies, and their consequent dermatopathological analysis, continue to be an important tool for the study of skin diseases and cutaneous manifestations of systemic diseases. Hematoxylin and eosin (H&E) staining constitutes the basic routine method for the microscopic observation of skin sections. In addition, trichrome stains are used for the staining of connective tissues and, initially, to discriminate between collagen and smooth muscle. Several trichrome stains are available and are preferentially applicable to specific connective tissues. So, staining of the skin with the classical and widely used Masson trichrome provides brilliant results and a clear discrimination between the epithelial (i.e., epidermis and derivatives) and mesenchymal (i.e., dermis) skin components (Fig. 15). In this sense, the RGB trichrome provides an equivalent differential staining of the epidermis and the dermis, but is particularly useful for objective quantification of collagen under polarized light microscopy (Fig.15)^**34**,**35**^.

**Figure 15.**
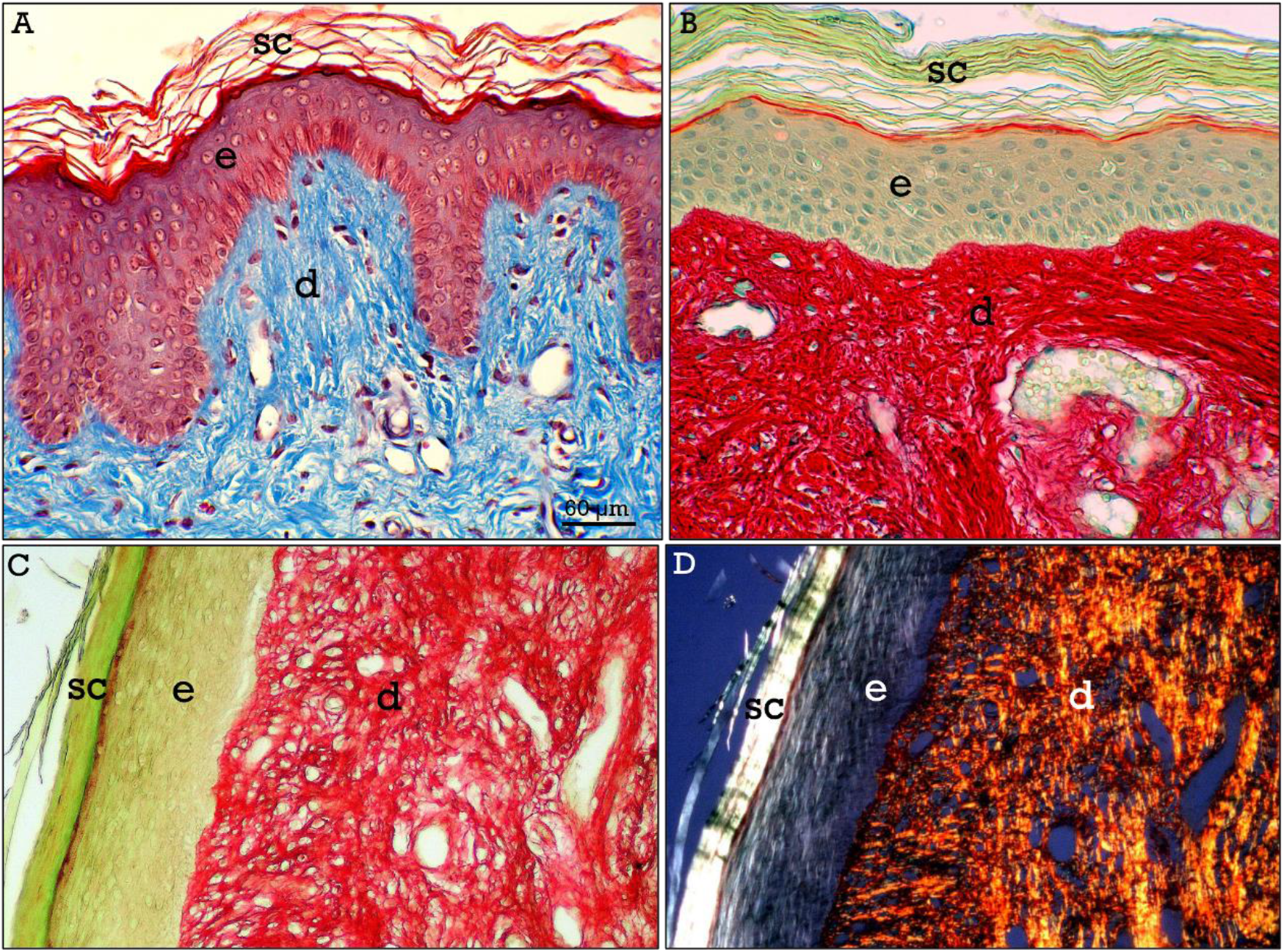
Comparison of Masson and RGB trichrome I. Staining with Masson trichrome (**A**) provides brilliant results and a clear contrast between the dermis (***d***) and the epidermis (***e***), similarly to RGB tricrome (**B**), although a clear advantage of the former is the staining of cell nuclei. However, RGB staining method allows objective collagen quantification when RGB stained sections (**C**) are observed under polarized light (**D**), as collagen show a specific red/orange birefringence.

Overall, the RGB trichrome provides a general staining of the skin extracellular matrix, and differential staining of the dermal components (i.e., nerves, secretory glands, etc), and is particularly useful for the study of dermal proteoglycan-enriched zones that cannot be clearly appreciated by using other common trichrome stains such as Masson trichrome, as the staining of the PEB is masked by the general staining of the dermal collagenous tissue (Fig. 16A,C), while the RGB trichrome produces a highly contrasted interface between the PEB (blue-stained) and the surrounding red-stained collagenous tissue (Fig. 16B,D).

**Figure 16.**
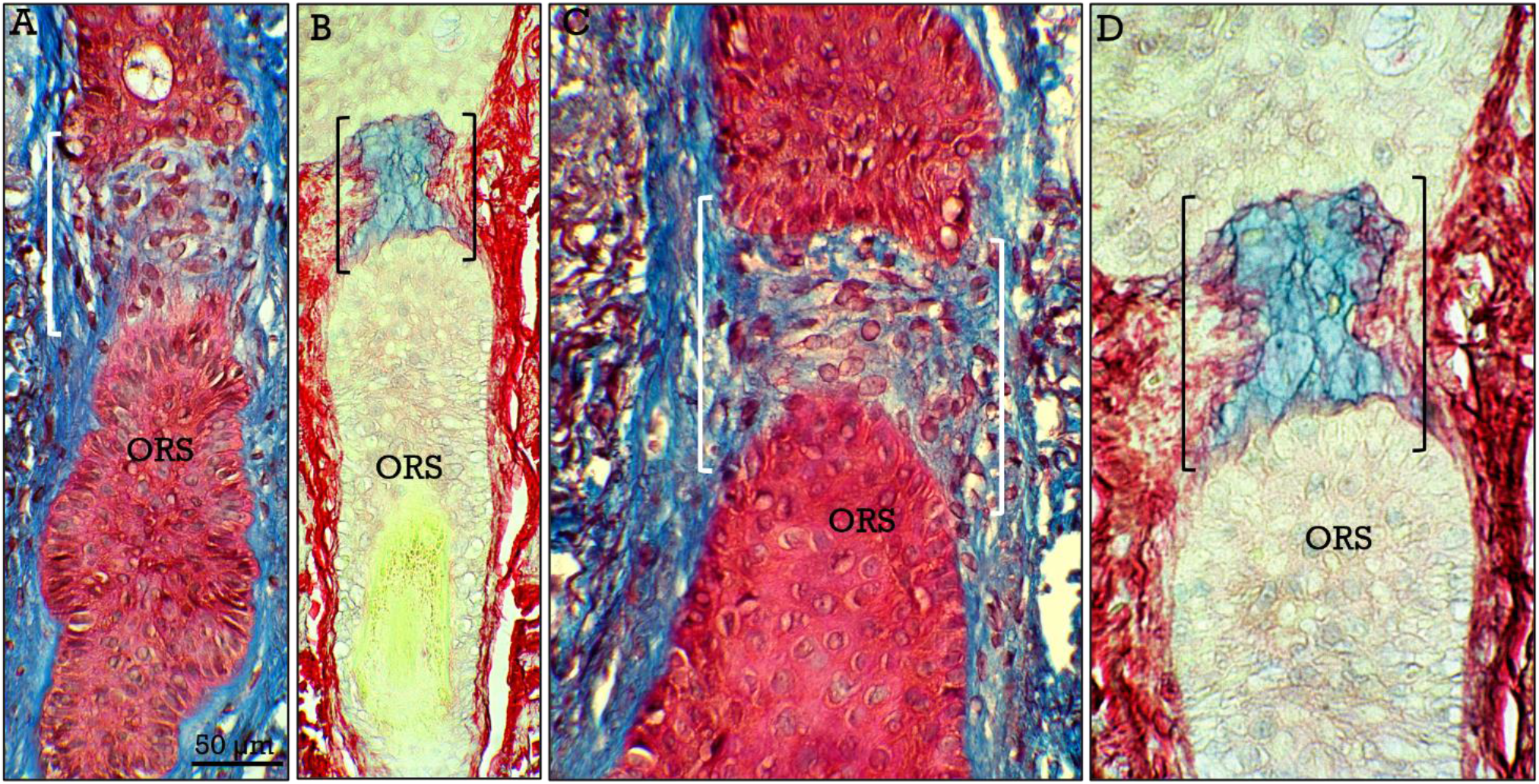
Comparison of Masson and RGB trichrome II. Areas containing abundant proteoglycans, such as the dermal papilla and the PEB (***brackets***), cannot be appreciated in follicles stained with Masson trichrome (**A**,**C**), in contrast to RGB-stained sections (**B**,**D**) in which collagen and proteoglycans show different (red *vs* blue) colors.

The RGB trichrome includes two primary dyes that stain two of the main components of the extracellular matrix in contrasted non-overlapping colors: sirius red, that specifically stains collagen, and alcian blue that stains glycosaminoglycans and proteoglycans. The resulting combined staining provides information about the relative amounts of collagen and proteoglycans in each particular connective tissue extracellular matrix. Intense alcian blue stained areas of the dermal sheath occur at two specific locations, the papilla located at the bulb, and the PEB located at the upper isthmus. Glycosaminoglycans/proteoglycans have been reported to play important roles in the interaction between epithelial cells and extracellular matrix, participating in the regulation of cell proliferation in different tissues^**37–39**^. Although proteoglycans are found thoughout the dermal sheath^**40**^, the prevalent presence of collagen determines its global red staining. However, the blue staining of the dermal papilla and the PEB strongly suggest that proteoglycans are particularly abundant in these areas, thus resulting in global alcian bue staining. Yet, the possible functional role of the PEB remains to be determined.

Several stem cell populations have been reported to be present at different zones of the hair follicle. The first described of these stem cell populations corresponds to that located at the attachment of the arrector pili muscle, known as the bulge area^**8**^. These stem cells express a number of specific markers such as Lgr5 and CD34 and are immersed in a niche composed of the extracellular matrix of the bulge, as well as the smooth muscle cells of the APM. An additional stem cell population has been described at the upper isthmus in the mouse hair follicles^**41**^, just below the junction of the hair follicle with the sebaceous gland duct. These stem cells express several stem cell markers such as Lgr6 and MTS24^**41**,**42**^, and could give rise to all skin cell lineages^**41**,**43**^. An equivalent population of Lgr6+ cells have been reported to be present in the same area of human hair follicles^**44**^. This location is coincident with that of PEB in the surrounding dermal sheath, thus suggesting that the PEB could be a crucial component of the niche for this stem cell population. Some studies have also related the expresion of Lgr6 marker to the presence of nerves at this zone^**45**^. Despite the enormous advances in the field of skin regeneration achieved during the last decades, the regulation of the different hair follicle stem cell populations is not fully understood and remains an enigma^**46**^. Anyway, the interactions between stem cells and extracellular matrix in the niche plays crucial roles in stem cell activation and differentiation. Hyaluronic acid and proteoglycans are fundamental components of the skin extracellular matrix and fulfill multiple functions^**47**^ as a component of the skin stem cell niche^**48**^. In this context, the proteoglycan-enriched area of the dermal sheath unveiled by RGB staining (PEB) deserves additional investigation.

In addition, RGB trichrome highlights the alterations of elastic fibers in sun-exposed skin, that is an almost physiological condition during skin aging in light-skinned people^**36**^. The color contrast between altered elastic and collagen fibers in RGB stained sections is particularly important at early stages of actinic elastosis (Fig. 17). In contrast, a clear general advantage of Masson trichrome is the staining of cell nuclei, an aspect in which the RGB trichrome gives poor results. For this reason RGB trichrome should be complemented with parallel H&E staining.

**Figure 17.**
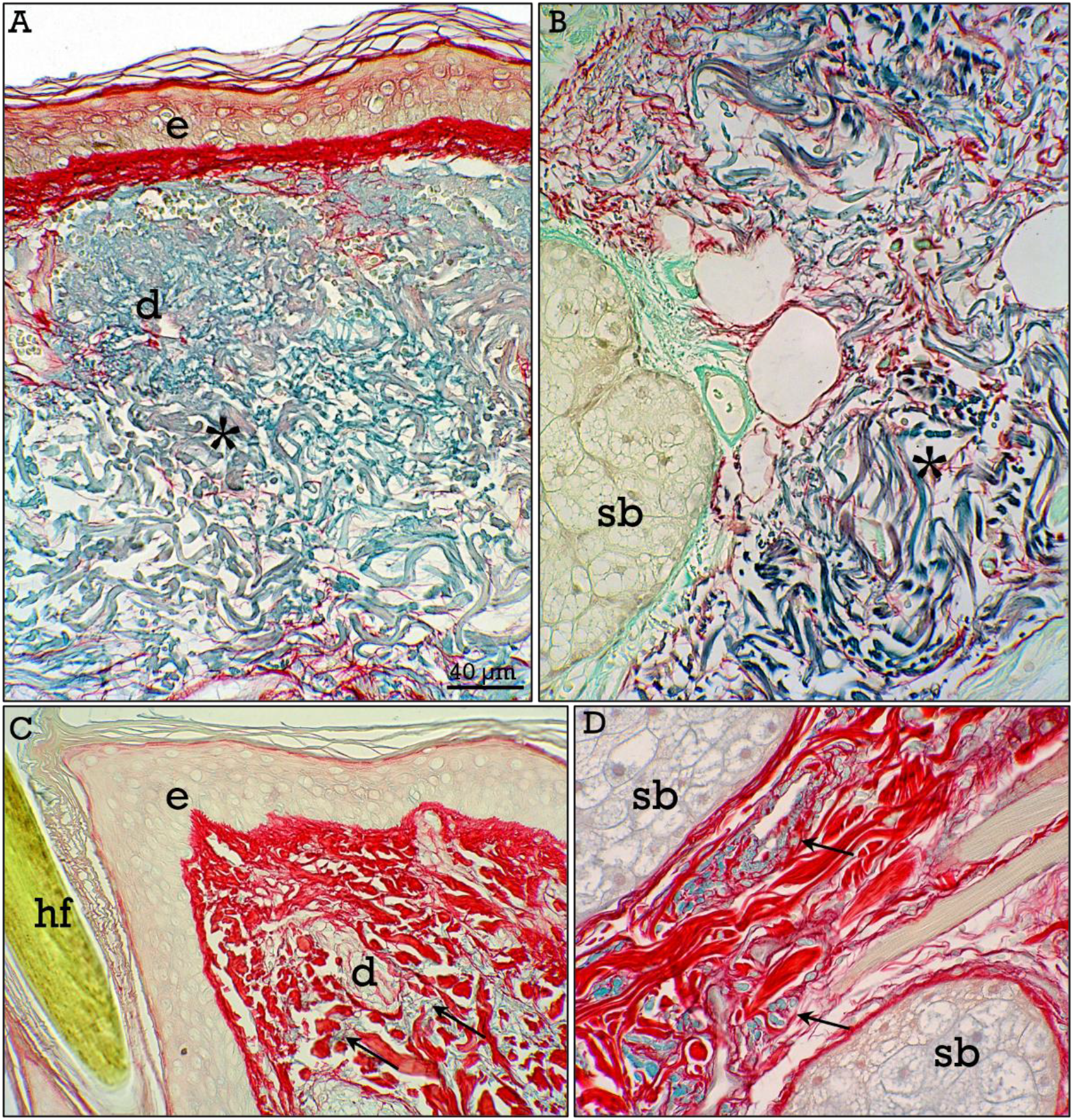
Staining of actinic elastosis with RGB trichrome. Actinic elastosis in the dermis (***asterisks***) is highlighted in RGB stained sections of sun-exposed aged skin (**A**,**B**), displaying a clear distinction between degenerated elastic (blue-stained) and remaining collagen (red-stained) fibers. This is particularly relevant at initial stages (**C**,**D**), when degenerated elastic fibers (***arrows***) are still scarce.

Summarizing, the RGB trichrome provides an useful staining method applicable to the human skin, that allows objective collagen quantification through polarized light microscopy, and reveals two domains of the dermal sheath that are particularly rich in proteoglycans.

## Acknowledgements

The authors are very grateful to María Cantillo and Esteban Tarradas for their technical assistance.

## Author contribution

Both authors were involved in conducting the study and contributed to the final version of the manuscript.

